# More than just oil droplets in water: surface tension and viscosity of protein condensates quantified by micropipette aspiration

**DOI:** 10.1101/2021.05.28.446248

**Authors:** Huan Wang, Fleurie M. Kelley, Dragomir Milovanovic, Benjamin S. Schuster, Zheng Shi

## Abstract

The material properties of biomolecular condensates play pivotal roles in many biological and pathological processes. Despite the rapid increase in the number of biomolecules identified that undergo liquid-liquid phase separation (LLPS), quantitative studies of the resulting condensates have been severely lagging behind. Here, we develop a micropipette-based technique, which uniquely allows quantifications of both the surface tension and viscosity of biomolecular condensates, independent of labeling and surface wetting effects. We demonstrate the accuracy and versatility of this technique by measuring condensates of LAF-1 RGG domains and a polymer-based aqueous two-phase system (ATPS). We anticipate this technique will be widely applicable to biomolecular condensates and will resolve several limitations regarding current approaches.

## Main Text

Biomolecular condensates that arise from LLPS have recently emerged as a central player in numerous cellular processes^1,2^. Surface tension and viscosity are two independent parameters that define the material properties of a liquid^3,4^. Gradual increases in the viscosities of biomolecular condensates are often linked to the formation of fibrils that underlie aging-associated diseases^5–10^. Quantification of condensate rheology therefore holds promise for unravelling the mechanisms, as well as facilitating therapeutic advances in the treatment of these diseases^11^.

While changes in condensate viscosity often have pathological consequences, the surface tension of biomolecular condensates can play key physiological roles: differences in surface tension can lead to layered multi-phase condensates, such as the compartmentation in nucleoli^12–14^. During autophagy, surface tension determines whether p62 condensates will be sequestered in small droplets or digested as a whole^15^. Finally, the nucleation of microtubule branches relies on an instability of TPX2 condensates, driven solely by the condensates’ surface tension^16^.

Several techniques have been developed to probe either the viscosity or the surface tension of biomolecular condensates^12,17–21^. The most widely used measure of viscosity relies on fluorescence recovery after photobleaching (FRAP), which is challenging to quantify in the scenario of 3-dimensional compartments such as biomolecular condensates^4,22^. Measurements of surface tension rely heavily on the fusion kinetics between two condenstates^17^. While significant improvements have been made^23,24^, the fusion assay is intrinsically limited because only a ratio of surface tension to viscosity can be estimated^12^. Therefore, a user-friendly technique that can directly measure both surface tension and viscosity of biomolecular condensates is still missing.

Micropipette aspiration (MPA) has been well-established to study the elastic properties of cells and liposomes ^25,26^. However, it has been challenging to apply MPA to quantify liquids. For MPA to perform well, the viscosity of the liquid needs to be large for the flow process to be adequately captured. At the same time, the surface tension of the liquid needs to be small for the aspiration pressure to overcome the capillary effect (see Methods). Both requirements are in contradiction to the properties of common liquids, where low viscosity (10^−3^ ~ 10^−2^ Pa·s) and high surface tension (~10 mN/m) are often observed. However, currently available data suggest that biomolecular condensates exhibit high viscosity (~10 Pa·s) and low surface tension (10^−3^ ~ 10^−2^ mN/m), making them uniquely poised for quantitative MPA studies (Figure 1).

**Figure 1.**
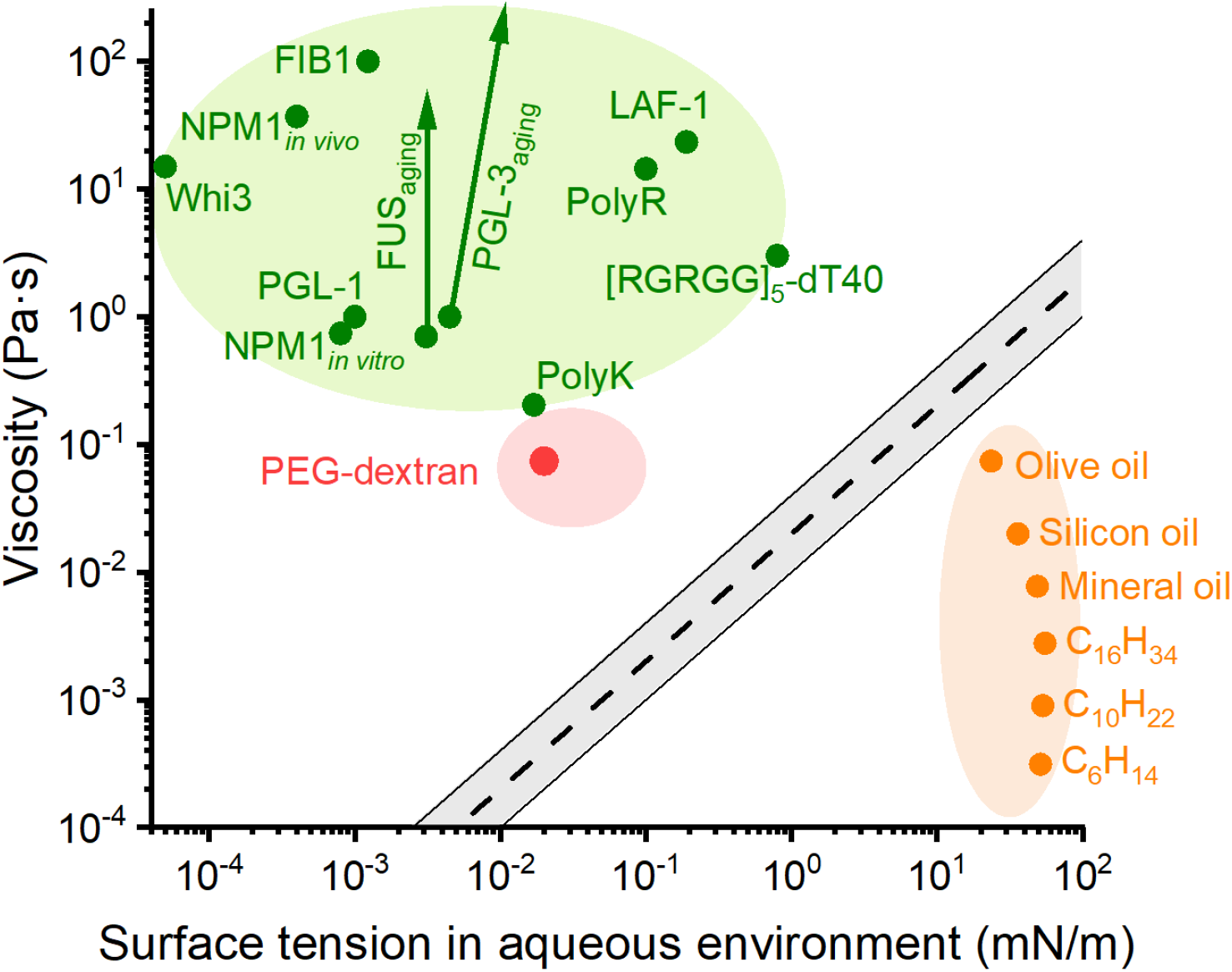
Viscosity and surface tension of liquids. Viscosity and surface tension of biomolecular condensates in aqueous buffer (green, arrows represent changes of properties over time) and common ‘oil droplets’ in water (orange). The gray belt represents an estimated boundary above which MPA will be well-suited for viscosity and surface tension measurements (Methods). The red and pink regions represent values for dextran-rich condensates measured in this study and estimated from literature^29,30^, respectively. See Table S1 for values and references used in this plot.

Here, we demonstrate the application of MPA to quantify the viscosity and surface tension of liquid condensates. We calibrated our method using a PEG-dextran ATPS^27^. This allowed us to develop a linear model to extract the material properties of condensates from their responses to MPA. We applied this technique to quantify condensates formed by the RGG domain, a well-known RNA binding region of the P granule RNA helicase LAF-1^18,28^. We further confirmed our viscosity and surface tension measurements by FRAP and fusion assays, respectively. Our results suggest that material properties of protein condensates are closer to ATPS than to oil droplets in water. MPA represents an active microrheology technique that can simultaneously quantify independent properties of biomolecular condensates, insensitive to common sources of artifacts such as labeling, photobleaching, and wetting effects of proteins.

To study the stress-strain relation of a liquid, a critical aspiration pressure *P_γ_*, determined by the surface tension (*γ*) of the condensate, needs to be reached. At aspiration pressures (*P*_asp_) greater than *P_γ_*, the condensate will flow into the micropipette (Figure 2a, we define suction pressures as positive). For a Newtonian fluid, the pressure difference and the flow rate are linearly related via the condensate’s viscosity (*η*):

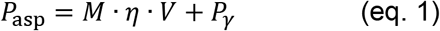

**Figure 2.**
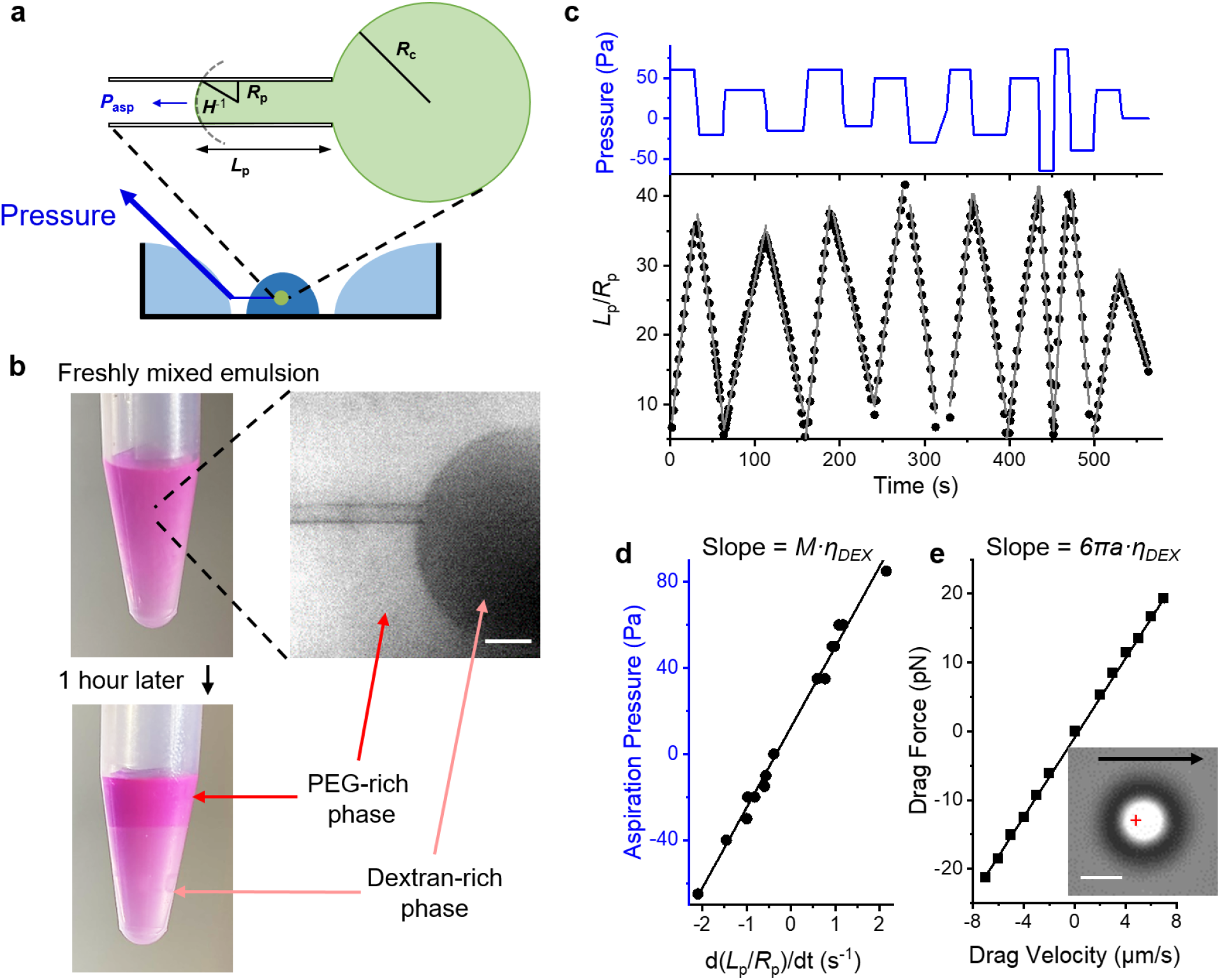
Calibration of MPA with PEG-dextran ATPS. **a**, Illustration of the micropipette aspiration system, dark blue: sample, light blue: water, green: a pipette-aspirated condensate, *P*_asp_: aspiration pressure, *R*_p_: pipette radius, *L*_p_: aspiration length, *R*_c_: radius of condensate outside the pipette. *H*: mean curvature of the liquid interface in the micropipette (positive direction as illustrated). **b**, An emulsion of PEG-dextran (upper) undergoes bulk LLPS after creaming for 1 hour (lower). Scale bar, 20 μm. **c**, Aspiration pressure (upper) and normalized aspiration length (lower) during MPA. Gray lines: linear fits of the normalized aspiration length for each pressure step. **d**, *P*_asp_ of each step plotted against *V* (slopes of the gray lines in **c**). **e**, Viscosity determination by optical dragging. Inset image: a trapped polystyrene particle dragged at 5 μm/s (arrow) in the dextran-rich phase. Cross: trap-center. Scale bar, 2 μm. Linear fits: *R*^2^ = 0.990 for **d** and 0.999 for **e.**

Here, *V* = *d*(*L*_p_/*R*_p_)/*dt* is the normalized flow rate. The critical pressure *P_γ_* = 2*γ*(*H* – 1/*R_c_*). *L*_p_, *R*_p_, *R*_c_, *H* describe the shape of the aspirated condensate and are readily available through microscopy (Figure 2a). However, the unitless factor *M*, which is a constant when (*R*_p_/*R*_c_)^3^ ≪ 1, must be determined experimentally by aspirating liquids of known viscosities^31^. Then, by measuring *V* under different *P*_asp_, the viscosity and surface tension of an unknown liquid condensate can be quantified from the slope and intercept, respectively, of eq. 1.

To calibrate ‘*M*’ with liquids that are appropriate for MPA (Figure 1), we chose an ATPS composed of PEG (8000 Da) and dextran (500,000 Da)^15,29,32^. Under a range of concentrations, mixtures of PEG and dextran will phase-separate into emulsions of micrometer-size droplets (Figure S1a). Rhodamine-B was included to identify the condensates microscopically. This mixture produced a labeled emulsion that is stable on the timescale of MPA experiments (~10 min), but undergoes bulk phase separation after 1~2 hours (Figure 2b).

Stepwise aspiration pressures were applied to dextran-rich condensates, and the aspiration length was found to change linearly under each pressure step (Figure 2c, Figure S1b). The resulting relation between the aspiration pressure and the condensate flow rate (Figure 2d), agrees well with predictions of a Newtonian fluid. The slope d*P*_asp_/d*V* (37.0 ± 0.7 Pa·s; n = 6; mean ± SEM for all values reported herein), therefore, represents the viscosity of the dextran-rich phase multiplied by ‘*M*’ (eq.1). We then directly measured the viscosity by dragging an optically-trapped particle within the dextran-rich phase (Figure 2e). The measured viscosity of the dextran-rich phase (74 ± 4 mPa·s) agreed with bulk viscometer measurements (Methods), giving *M* = 500 ± 30. Additionally, the intercept from the *P*_asp_-*V* relation corresponds to a surface tension of 0.02 ± 0.01 mN/m (eq. 1), in agreement with the literature^29^. With careful control of water evaporation (Figure S2), next we applied MPA to protein samples of limited volumes (20~30 μL).

LAF-1 is one of the first well-studied proteins that undergo LLPS, mainly due to its intrinsically disordered N-terminal RGG domain^4,18,19,22,28^. We applied MPA to an engineered tandem RGG domain that robustly undergoes LLPS (Methods; hereafter named RGG condensates)^28^. Unlike dextran, RGG condensates fully wet the inner wall of the micropipette, requiring a negative *P*_γ_ to balance the capillary effect (Figure 3a, *H* ≈ −1/*R*_p_). Beyond *P*_γ_, the aspiration length changed linearly under each pressure step (Figure 3b, Movie S1), indicating a lack of condensate elasticity at the timescale we were probing (>1 s). After initial-entry steps, *V* increased linearly with *P*_asp_ (Figure 3c, S3, S4). The slope corresponded to a viscosity of 1.62 ± 0.18 Pa·s (n = 11), in agreement with previous estimates^33^. As expected from the wetting behavior, the intercept of *P*_asp_ vs. *V* was negative, and the corresponding surface tension was 0.159 ± 0.010 mN/m (n = 11).

**Figure 3.**
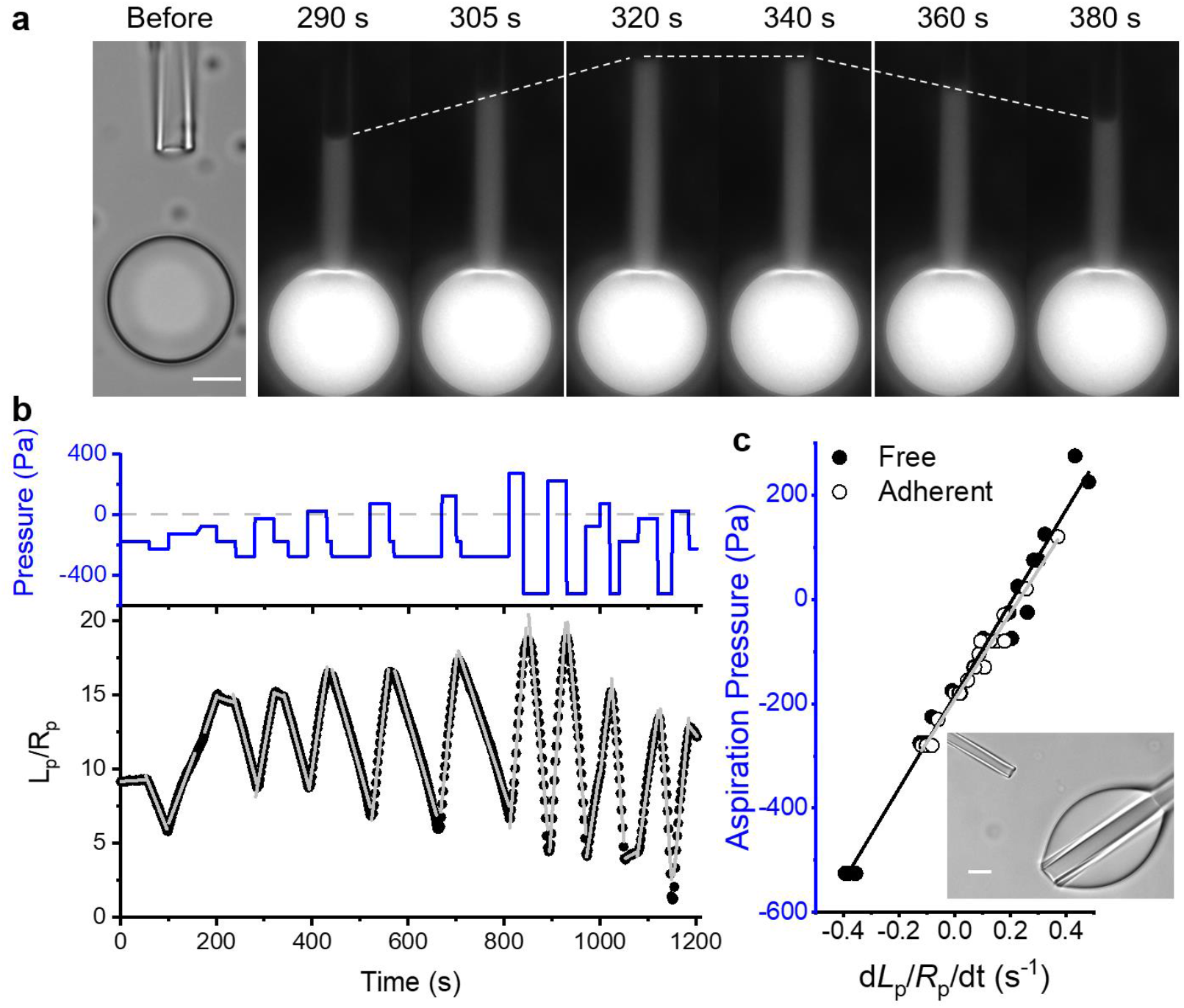
MPA of RGG condensates. **a**, Left: transmitted light image of an RGG condensate with a nearby micropipette. Right: time lapse fluorescent images of the RGG condensate under three aspiration pressures: −25 Pa (290~320 s), −175 Pa (320~340 s), and −275 Pa (340~380 s). Dashed lines trace the change of *L*_p_. **b**, Aspiration pressure (upper) and normalized aspiration length (lower) during MPA. Dashed line: zero pressure. Gray lines: linear fits of the normalized aspiration length for each pressure step. **c**, *P*_asp_ of each step plotted against *V* for a free condensate (closed, see **a**) and a condensate strongly adhered to a glass pipette (open, inset image, see Methods). Linear fits to the data are shown as black (slope: 890 ± 30 Pa·s, intercept: −184 ± 7 Pa, R^2^ = 0.982) and gray (slope: 830 ± 30 Pa·s, intercept: −192 ± 4 Pa, R^2^ = 0.976) lines, respectively. All scale bars, 10 μm.

Many proteins tend to wet and adsorb onto solid surfaces^4,16,23,24,34^. While potentially mediating important biological processes^15,16,34^, this wetting effect can introduce significant artifacts in measurements that rely on the fusion kinetics or morphology of condensates^12,23,24^. In contrast, the contribution of *R*c in eq. 1 is negligible when *R*c^−1^≪|*H*|, making MPA insensitive to the wetting of condensates (Figure 3c, Movie S2). Additionally, fluorescent labeling of the protein is not necessary for MPA as long as the condensate-buffer interfaces can be resolved (Figure S1c, S4, and Movie S3). For the same reasons, MPA measurements are insensitive to photobleaching and can be easily combined with fluorescence-based studies^24,35^, further expanding the applicability of this technique.

To confirm the surface tension and viscosity of RGG condensates measured by MPA, we first adopted an improved version of the condensate fusion assay^23,24^. Two optically-trapped RGG condensates were manipulated to encounter and the subsequent fusion process was recorded (Figure 4a, 4b). A linear relation was observed between the fusion time and the size of the condensates (Figure 4c). The slope, which scales with the inverse capillary velocity *η/γ*, was 0.016 ± 0.002 s/μm, in agreement with the MPA measurements (*η/γ* = 0.010 ± 0.001 s/μm). We then used FRAP to estimate the viscosity of RGG condensates. A circular region within RGG condensates was photobleached, and diffusion coefficients were calculated based on the half-recovery time (Figure 4d)^22^. Combined with an estimate of the protein hydrodynamic radius, we obtained a viscosity of 1.8 Pa·s, comparable to the MPA result. We noticed that viscosity values between 0.8 and 3.6 Pa·s can be extracted from FRAP, depending on the extract model of choice (Methods, Figure 4e).

**Figure 4.**
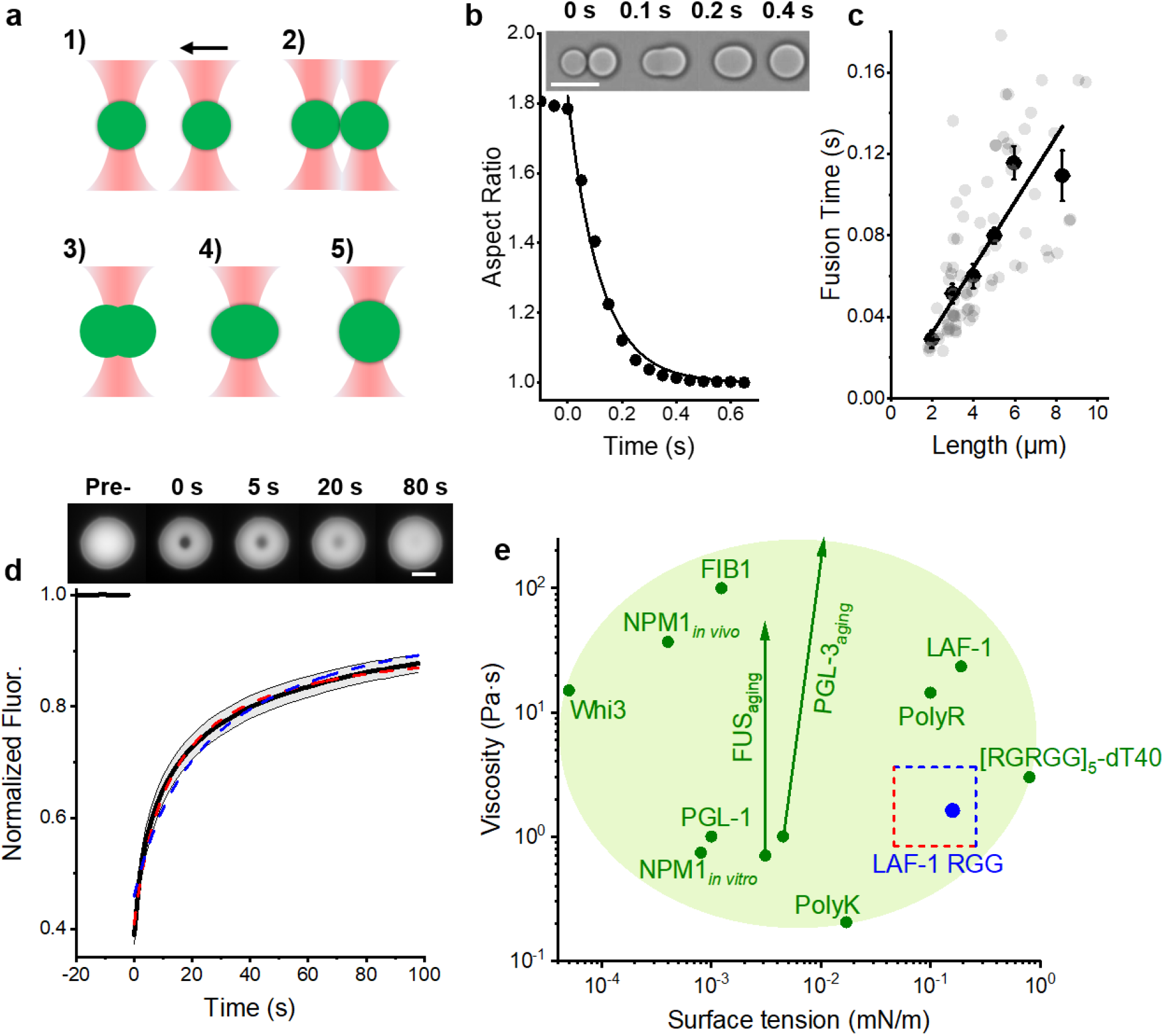
Surface tension and viscosity of RGG condensates estimated from condensate fusion and FRAP. **a**, 1)-5): Illustration of the condensate fusion experiment using dual-optical traps. **b,** Fusion of two RGG condensates (inset images) quantified as a decrease of the overall aspect ratio to 1. The curve is an exponential fit. **c,** The fusion time of RGG condensates vs. the condensates’ length. Gray and black circles are individual (n = 76 pairs) and binned fusion experiments, respectively. Line: weighted linear fit to the binned data (slope = 0.016 ± 0.002 s/μm, R^2^ = 0.902). **d**, FRAP measurements within RGG condensates (n = 42). Inset images show a representative experiment. Red dash: fit to 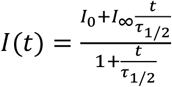, with *τ*_1/2_ = 12.4 ± 0.5 s and *I*_∞_ = 0.928 ± 0.003, R^2^ = 0.995. Blue dash: fit to 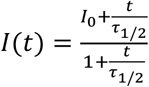, with *τ*_1/2_ = 24 ± 1 s, R^2^ = 0.966. **e**, Zoom-in of Figure 1, with viscosity and surface tension of RGG condensates measured by MPA in blue circle. Estimates from FRAP and fusion kinetics are represented by the dished box (Methods). All error bars are SEM, all scale bars are 10 μm.

RGG condensates fuse quickly (~0.1 s) with >90% of the constituting proteins moving freely, consistent with their liquid behavior during MPA. However, many biomolecular condensates can take more than 100 s to fuse^15,36^, while exhibiting small fractions (<50%) of mobile proteins^3,10,37,38^. In the latter cases, MPA measurements will be essential to clarify confusion around the condensates’ material properties^10^. The unambiguous quantification of RGG condensates through MPA further iterates the contrasting material properties of biomolecular condensates and oil droplets (Figure 1). For example, mineral oil has a surface tension 300-fold higher than that of RGG condensates, whereas its viscosity is more than 200-fold lower.

Surface tension and viscosity are tunable through intermolecular interactions. Thus, a future direction will be to systematically dissect how protein sequence and biochemical environment affect the material properties of biomolecular condensates^4^. Importantly, by implementing a whole-cell patch-clamp configuration^39^, MPA can be applied to study biomolecular condensates *in vivo*. Finally, MPA setups are readily available in electrophysiology and biomechanics labs, making it easily adaptable for studying the material properties of biomolecular condensates in the broader biological and chemical communities.

## Acknowledgements

We want to thank Ehsan Atefi, Ningwei Li, Frank Jülicher, and Rick Remsing for helpful discussions. We thank Gayatri Ganeshan, Steven Arnold, and Roberto Sul for critical comments on the manuscript. We also thank Andy Nieuwkoop and his lab for helping with materials and storage space. Z.S. and B.S.S are supported by Rutgers University startup funds. F.M.K is supported by NIH Award T32 GM135141. D.M. is supported by the German Research Foundation (DFG MI 2104 and SFB 1286) and the German Academic Exchange Service (DAAD PPE 2021).

## Supplementary information includes

## Methods

### Estimate of the working range of micropipette aspiration (MPA) through dimensional analysis

During micropipette aspiration of a liquid (Figure 2a), the viscosity needs to be large in order for the camera to capture the flow process. At the same time, the surface tension needs to be small in order for the flow to start.

Assume the maximal imaging frequency is 100 Hz (*Δt*_min_ = 0.01 s), the radius of the pipette: *R*_p_ = 1 μm, and *M* = 500 (see eq. 1 in the main text). In order to capture liquid deformations that are on the order of pipette diameter (*ΔL*_p_ = 2 μm), the viscosity *η* (in Pa·s) needs to satisfy:

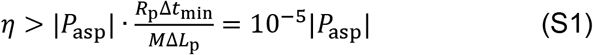

The aspiration pressure needs to overcome the capillary effect caused by the surface tension *γ* (in mN/m). For a non-wetting (*H*^−1^ = *R*_p_) or a perfectly wetting (*H*^−1^ = −*R*_p_) liquid with *R*_c_ ≫ *R*_p_:

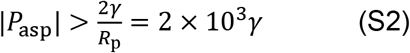

Combine relations S1 and S2:

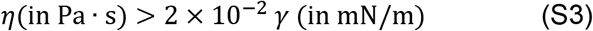

Therefore, relation S3 defines the regime of viscosity and surface tension where MPA is expected to perform well. In Figure 1, *η* = 0.02 *γ* is plotted as the black dashed line, gray regions represent *η* = 0.01 *γ* ~ 0.04 *γ*.

### Protein purification and sample preparation

RGG-based proteins were expressed recombinantly in *E. coli* and purified by affinity chromatography, as previously described^1^. The working protein sample contains 1 μM RGG-EGFP-RGG (molecular mass 62.1 kDa) and 6 μM RGG-RGG (molecular mass 35.7 kDa) in a pH 7.5 buffer containing 20 mM Tris and 150 mM NaCl.

Phase separated dextran and PEG aqueous two-phase systems were prepared by mixing different concentrations of PEG-8000 (43443-22, Alfa Aesar, US) and dextran-500k (DE132-100GM, Spectrum Chemical, US) stock solutions. The stock solutions were prepared by dissolving each polymer in Milli-Q water. Emulsions of different PEG to dextran ratios showed different distributions of droplet size (Figure S1a). The 5% PEG and 6.4% dextran (both by mass) mixture was chosen for micropipette aspiration, because the resulting emulsion contained droplets with comparable sizes to those of the protein condensates.

Rhodamine-B (83689-1G, Sigma, USA) was added (at a final concentration of 1 μM) to the PEG-dextran mixture to distinguish the dextran phase from the PEG phase (Figure 2b). Rhodamine-B preferentially enters the PEG-rich phase^2^, therefore dextran-rich condensates showed as dark droplets in a bright background in fluorescent microscopy images (Figure 2b, upper right image). The fluorescent labeling was confirmed by the observation that after bulk LLPS, the heavier dextran-rich layer (Figure 2b, lower layer of the lower left image) contained less Rhodamine-B compared to the lighter PEG-rich layer. The concentration of dextran in the dextran-rich layer was estimated to be ~14% by mass.

### Micropipette fabrication, aspiration, and imaging

Micropipettes were pulled from glass capillaries using a pipette puller (PUL-1000, World Precision Instruments (WPI), US). The tip of the pipette was cut to an opening diameter between 1~ 5 μm and bent to ~40° using a microforge (DMF1000, WPI).

Micropipette aspiration and imaging were carried out on a Ti2-A inverted fluorescent microscope (Nikon, Japan) equipped with a motorized stage and two motorized 4-axes micromanipulators (PatchPro-5000, Scientifica, UK). A micropipette was filled with the same buffer as the protein (20 mM Tris and 150 mM NaCl, pH 7.5) using a MICROFIL needle (WPI) and subsequently mounted onto a micromanipulator. The rear end of the pipette was connected to an adjustable water reservoir. The pipette holder was then rotated so that the bent tip of the micropipette was parallel to the imaging plane. The aspiration pressure within the micropipette was controlled and recorded by adjusting the water level in the reservoir using a set of 5 ml, 20 ml, 50 ml, and 150 ml syringes connected to the reservoir.

The zero pressure of the system was calibrated before each MPA experiment, using a dilute solution of fluorescent nanoparticles. The zero pressure (*P*_0_) was set according to the point where fluorescent nanoparticles underwent Brownian motion inside the micropipette. The error in aspiration pressure (<2 Pa) was defined as the minimal pressure change near *P*_0_ that resulted in an observable directed flow of fluorescent particles in the micropipette.

MPA experiments were carried out in glass-bottom dishes (ES56291, Azer Scientific, US) that were pre-treated with 5% Pluronic F127 (P2443-250G, Sigma) for > 1 hour to prevent adhesion of RGG condensates to the glass^1^. Milli-Q water was added to the edge of the dish to minimize evaporation from the sample (Figure 2a). We further quantified water evaporation rates under our experimental conditions using a 20 μL sample of Rhodamine-B solution (Figure S2). Volume of the sample was assumed to be inversely proportional to its mean fluorescence intensity. No measurable volume change was observed when the dish-cap was on, providing a stable environment for necessary incubation periods for the sample. When the dish-cap was removed for micropipette aspiration, evaporation led to a slow constant decrease in the sample volume (~0.04 μL/min). We found that the evaporation can be compensated to be less than 5% in our MPA experiments (Figure S2).

After calibration of the aspiration pressure, a 20~30 μL sample of phase separated protein solution was added to the center of the dish (Figure 2a). Once micrometer-size protein condensates were observed at the bottom of the dish, a calibrated micropipette was moved to a condensate of interest to start the aspiration measurements. First, a positive (suction) pressure was applied to initiate the flow of the condensate into the micropipette. The condensate was typically allowed to flow into the micropipette until the aspiration length reached ~40 μm (the maximal aspiration length was limited by the field of view of the camera, initial condensate size, and the exact angle of the micropipette tip). Then, sequential stepwise ejection and suction pressures were applied to deform the condensate at different shear stresses while maintaining the aspiration length to be between 5 to 40 μm (Figure 3b and S3a-b). The deformation of the condensate was recorded using a 60X objective, at 1 Hz (ORCA-Flash 4.0, Hamamatsu, Japan), either through transmitted light imaging (Figure S1, S4, Movie S3) or through imaging the fluorescence of the EGFP tag (Figure 3, S3, Movies S1, S2). Of note, larger condensates typically result in more accurate MPA measurements, mainly due to the smaller perturbation of the changing aspiration length to *R*_c_. For a >10 μm condensate, typical changes in aspiration length correspond to a < 3% change in *R*_c_. Therefore, in our experiments, small condensates were first manipulated into a large condensate through either a micropipette or an optical trap (see “Optical trap mediated condensate fusion” section) before MPA measurements.

When the RGG condensate first entered the micropipette, wetting between protein and glass led to dramatic changes in the interfacial curvature between the condensate and buffer inside the micropipette. The interfacial curvature stabilized in later steps (Figure S3a, Movie S1). As a result, the *P*_asp_ vs. *V* relation during the initial-entry largely deviated from that of the remaining steps (Figure S3b, S3c). We corrected for the change in *H* by subtracting a time-dependent *P*_γ_ from the aspiration pressure (Figure S3d). However, the irreversible binding of a trace amount of protein to the inner wall of the aspiration pipette significantly accelerated the deformation of condensates during the initial-entry steps (Figure S3e, S3f). To account for the lack of information about the kinetics of protein-glass binding, we disregarded the measurements from the initial-entry steps.

After the initial steps, the interfacial curvature between RGG condensates and buffer in the micropipette was set by the wetting of the protein to the inner pipette wall (Movie S1). Due to this wetting effect, RGG condensates flowed into the micropipette under both positive (suction) and small negative (ejection) pressures, whereas decreases in aspiration length only happened under large negative (ejection) pressures (Figure 3, Movie S1, Figure S4).

Adhesion between RGG condensates and the glass bottom dish can be prevented by coating the glass with 5% Pluronic F127^1^. However, we noticed that the same coating procedure does not necessarily work for other protein condensates or glass surfaces. To expand the applicability of MPA, we compared the measurements on near-free condensates that sedimented to the bottom of coated dishes (Figure 3a) with condensates that are purposefully adhered to a bare glass pipette (Figure 3c). MPA gave near identical *P*_asp_ vs. *V* relations for the two types of RGG condensates (Figure 3c), even though *R*_c_ was not accurately defined in the strongly-adhered condensates. This agrees with expectations from eq. 1, where the contribution of *R*_c_ can be neglected when |*R*c^−1^|≪ *H*, as is in our experiments.

### Viscosity of dextran-rich condensates

To calibrate the viscosity measurements, MPA should be applied to condensates with viscosity values that can be easily determined through other means. Dextran-rich condensates in a PEG-dextran aqueous two-phase system were chosen for this purpose (Figure 2). After MPA, two independent methods were used to measure the viscosity of the dextran-rich phase.

1. Optical dragging An optical trap (Tweez305, Aresis, Slovenia) was applied to drag an *r* = 1.60 μm radius polystyrene bead (HUP-30-5, Spherotech, US) in a large dextran-rich condensate at 13 different velocities (Figure 2e). The slope of the dragging force (*f*) vs. dragging speed (*v*) was used to calculate the viscosity (*η*) based on the Stokes equation (eq. S4):

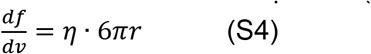 The measured viscosity was 74 ± 4 mPa·s. The stiffness of the optical trap (~ 0.02 pN/nm) was calibrated before each experiment by applying equipartition theorem to the thermal fluctuation of a trapped bead in the dextran-rich phase^3^.
2. Ubbelohde viscometer After the bulk-phase separation of 40 ml PEG-dextran mixture, the bottom layer, corresponding to the dextran-rich phase, was applied through an Ubbelohde viscometer (13-614C, Cannon Instrument, US). The viscosity was measured to be 80 mPa·s.

### Optical trap mediated condensate fusion

Two RGG condensates were individually controlled by two independent optical traps (Tweez305, Aresis, Slovenia) equipped on the Ti2-A inverted microscope (Nikon, Japan). As illustrated in Figure 4a, the right condensate was moved towards the left one until they touched. Then, the right optical trap was turned off, and the condensates were allowed to fuse under the combined influence of their viscosity and surface tension. The fusion processes were acquired at a frame rate of 20 Hz using a 60x water objective. The acquired images were analyzed in MATLAB (R2019a). The images were fitted into a Gaussian ellipse and the ratio of the major to minor axes of the ellipse (aspect ratio) was plotted as a function of time. The fusion time (τ) was extracted by fitting the change in the aspect ratio (*AR*) of fusing condensates to a single exponential decay (Figure 4b, eq. S5).

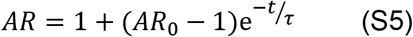

The length of condensates was defined as the geometric mean of the condensate diameters before fusion^4^. The ratio of viscosity to surface tension (inverse capillary velocity) was estimated from the slope of the fusion time vs. length relation (Figure 4c).

### FRAP measurement of the condensate viscosity

FRAP experiments were performed on a total internal reflection fluorescence microscope (DMi8 TIRF, Leica, Germany) equipped with an Infinity Scanner system (Leica, Germany). All images were acquired using a 100X oil objective at 1 Hz. A 1.5 μm radius circular region was photobleached at the center of large RGG condensates (radius 9 ± 2 μm) using a short pulse (~1 s) of focused 488 nm laser, and the fluorescence recovery was analyzed using ImageJ. After background subtraction, fluorescence of the bleached region (*I*_ROI_) was divided by the fluorescence of the entire condensate (*I*_cond_) according to eq. S6, to minimize photobleaching and boundary effects^5,6^.

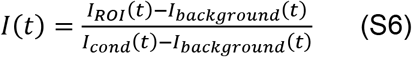

The time point right after the bleaching step was defined as time zero. *I*(*t*) was normalized so that the average of *I*(*t* < 0) equals to 1.

To extract the half-recovery time, *I*(*t*) was fitted to eq. S7a or eq. S7b, depending on whether an immobile fraction was included in the model.

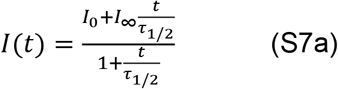

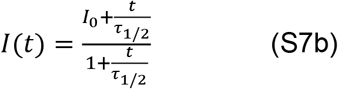

Here, *τ*_1/2_ is the half-recovery time and *I*_∞_ is the mobile fraction (in eq. S7b, *I*_∞_ is set to 1). See Figure 4d for the fitting results.

The diffusion coefficient (*D*) of the bleached molecule (RGG-EGFP-RGG) can be determined from a 2D or a 3D infinity model, according to eq. S8a or S8b, respectively^5^.

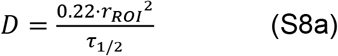

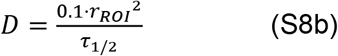

Where *r*_ROI_ = 1.5 μm is the radius of the bleached area, *τ*_1/2_ is the recovery time from eq. S7.

The viscosity of RGG condensates was then calculated using the Stokes-Einstein relation (eq. S9)^7^.

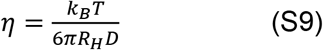

*R*_H_ is the hydrodynamic radius of RGG-EGFP-RGG. Using the online Hydrodynamic Radius Converter (https://www.fluidic.com/resources/Toolkit/hydrodynamic-radius-Converter/), *R*_H_ was estimated to be 6.54 nm, by taking into consideration the molecular mass and folding of RGG-GFP-RGG^8^.

In Figure 4e, the horizontal red dashed line (*η* = 0.8 Pa·s) represents viscosity calculated assuming the presence of immobile proteins (eq. S7a) in the 2D model (eq. S8a). The vertical red dashed line represents surface tension calculated using *η* = 0.8 Pa·s and the higher bound of inverse capillary velocity (*η/γ* = 0.018 s/μm) measured from the fusion assay (Figure 4c). The horizontal blue dashed line (*η* = 3.6 Pa·s) represents viscosity calculated assuming the absence of immobile fraction (eq. S7b) in the 3D model (eq. S8b). The vertical blue dashed line represents surface tension calculated using *η* = 3.6 Pa·s and the lower bound of inverse capillary velocity (*η/γ* = 0.014 s/μm) measured from the fusion assay (Figure 4c).

## Supplementary Figures

**Figure S1:**
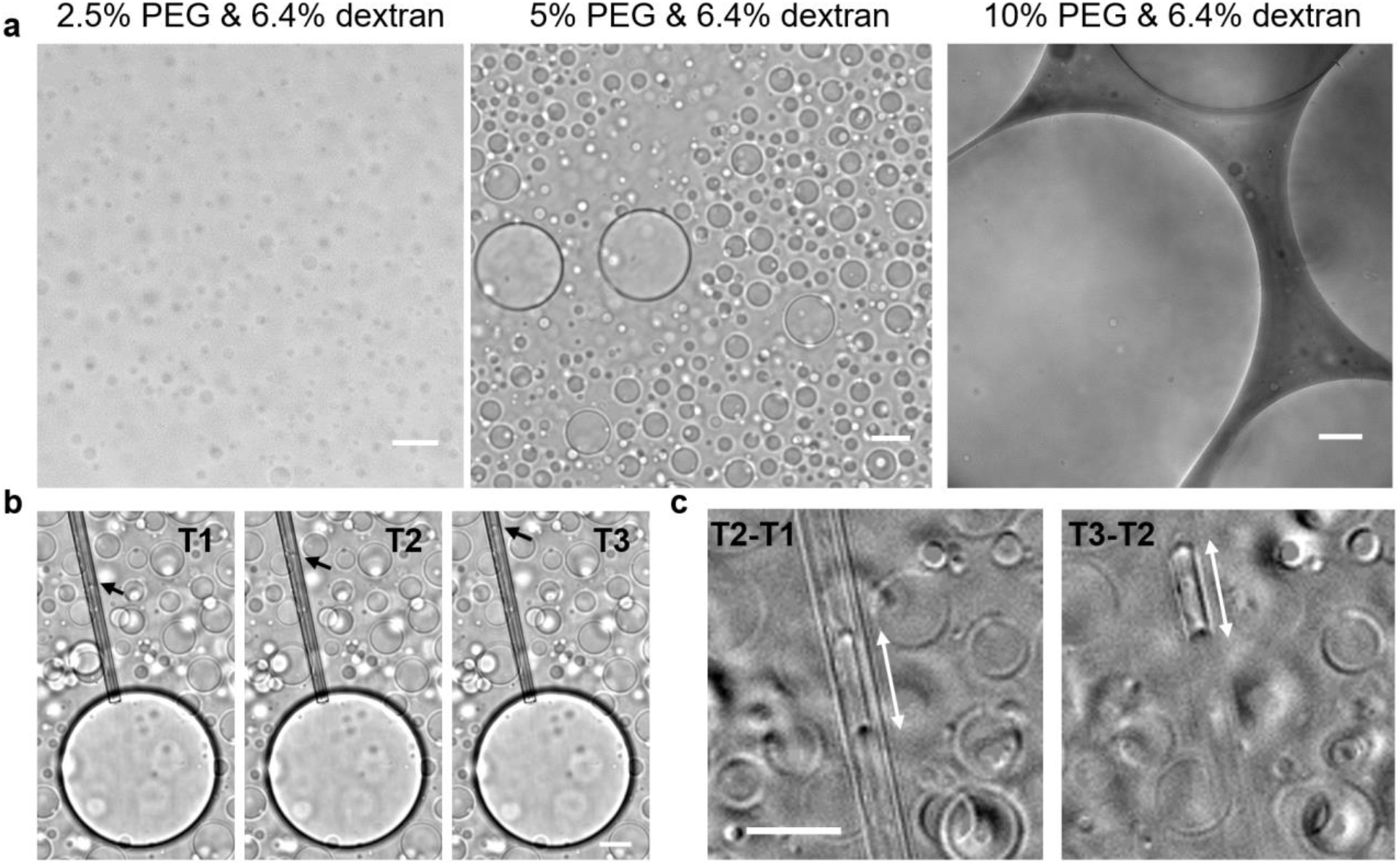
Phase separation and micropipette aspiration analysis of PEG and Dextran mixtures. **a**, Micrometer-scale droplets were observed in emulsions of PEG-dextran. Left to right: mixtures of PEG (8,000 Da) and dextran (500,000 Da) at increasing ratios of PEG to dextran. The 5% PEG & 6.4% dextran condition was chosen to produce droplets with similar sizes to those of protein condensates. **b**, Flow of a dextran-rich condensate into a micropipette (pre-filled with PEG-rich solution) under constant suction pressure (60 Pa). The 3 images were taken at 3 seconds apart. Arrows point to the interfaces between the dextran-rich and PEG-rich phases which are zoomed-in in **c**. **c**, Intensity differences between images in **b**: T2-T1 (left) and T3-T2 (right). The double-arrows show the increase of aspiration length in 3 seconds. Analysis of the MPA experiment can be achieved as long as the condensate-buffer interface is resolvable. All scale bars, 20 μm.

**Figure S2:**
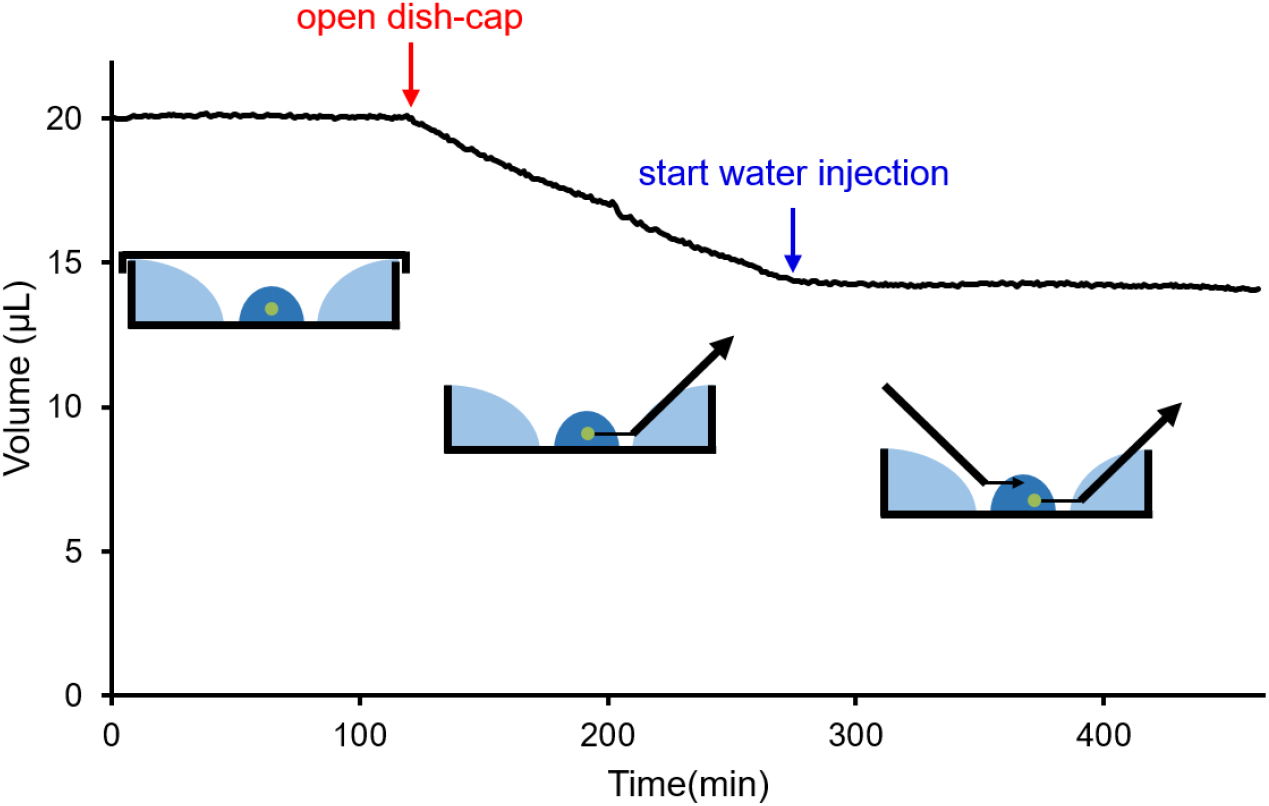
Quantification and correction of water evaporation during micropipette aspiration experiments. Under our experimental conditions, the presence of peripheral water eliminated evaporation from the 20 μL sample as long as the cap of sample dish was on. Upon removing the cap (red arrow) for micropipette aspiration, water slowly evaporated at a rate of 0.04 μL/min. The evaporation during micropipette aspiration was compensated (blue arrow) through continuous injection of pure water using a second micropipette, or by adding 2 μL of pure water every 50 min. Sample volume was measured through fluorescence-based concentration measurement of Rhodamine-B at an imaging rate of 1 frame per minute and no measurable photobleaching was observed.

**Figure S3:**
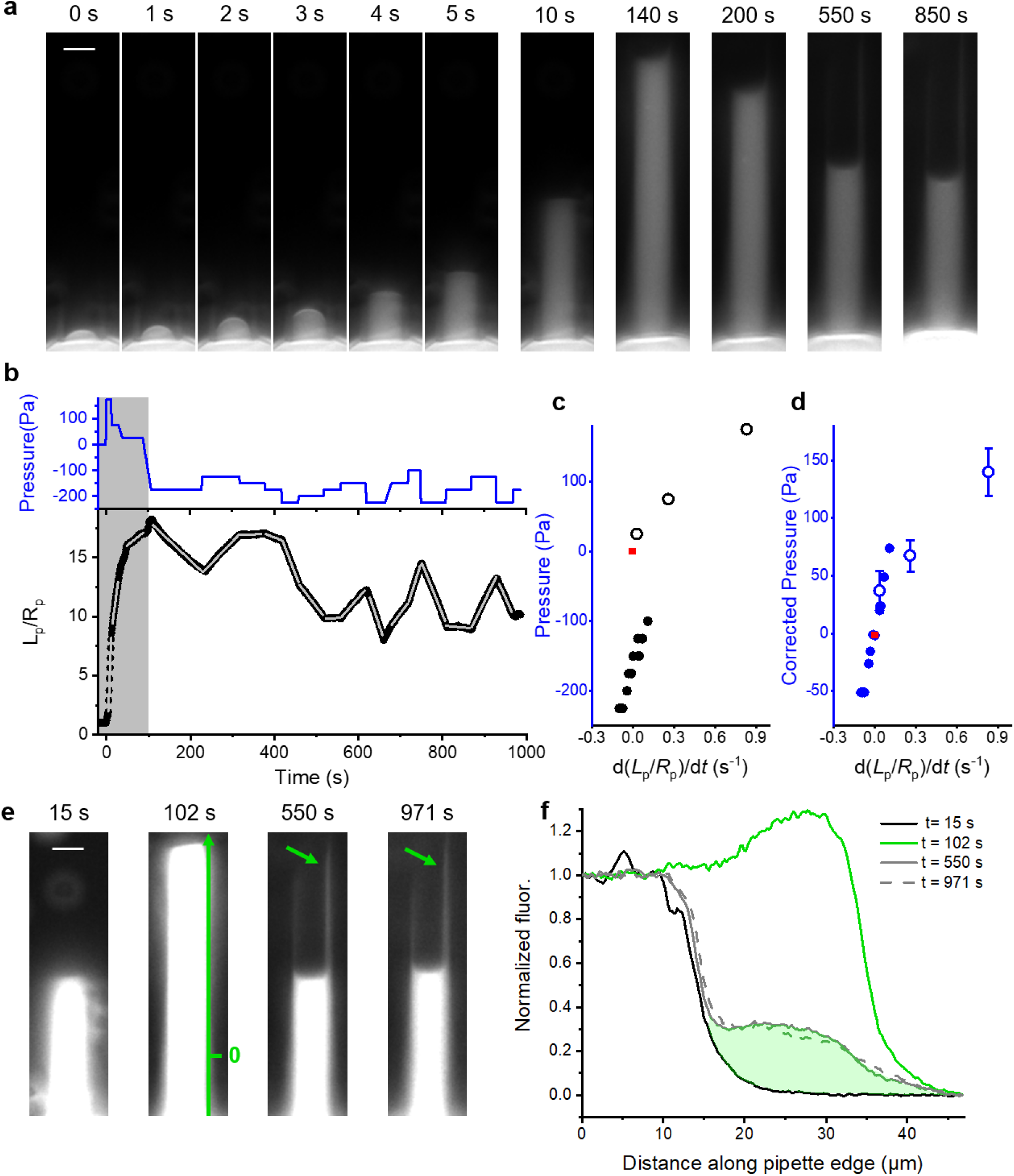
The interfacial curvature and the wetting of RGG condensate inside micropipette. **a**, Time lapse fluorescence images showing the aspirated portion of the condensate. After proteins enter the micropipette (1-4 s), the wetting of proteins to the inner pipette wall led to swift changes in the interfacial curvature of the protein condensate. In the case of RGG, this curvature stabilized within 2 min and remained near −1/*R*_p_ in the following aspiration steps. **b**, Aspiration pressure (upper) and normalized aspiration length (lower) as a function of time. Shaded area represents the initial-entry steps (defined as when the protein condensate first encountered a bare glass micropipette), where irreversible binding of protein to pipette inner wall happens. Gray lines are linear fits to the normalized aspiration length under each pressure step. **c-d**, Raw aspiration pressure (**c**) and tension-corrected pressure (**d**) of each step plotted against the normalized deformation rate (slopes of the gray lines in **b**). The initial-entry steps are denoted by open circles and a red square is placed at (0,0). Error bars in **d** reflect the uncertainty in interfacial curvature during the initial-entry steps. **e**, Over-exposed images of the aspirated portion during (15 s, 102 s) and after (550 s, 971 s) the initial-entry steps. The axis (at 102 s) represents the edge of the micropipette, arrow (at 550 s) points to proteins that were stuck to the inner pipette wall, which persisted in further aspiration steps (arrow at 971 s). **f**, Line profile along the pipette edge at the four time points shown in **e**. Area of the shaded region shows the amount of protein that was stuck on the inner wall of the micropipette. All scale bars, 5 μm.

**Figure S4:**
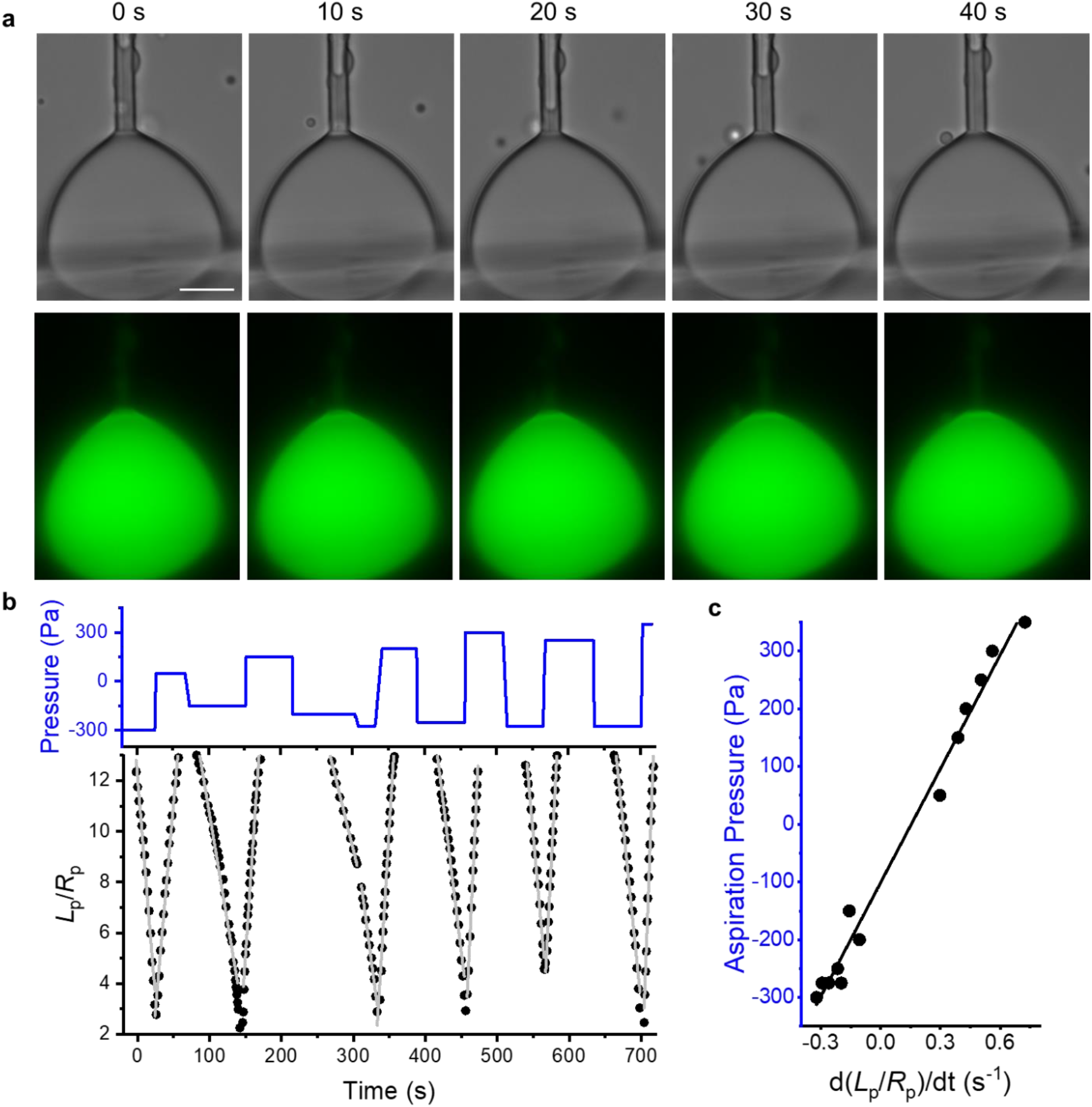
Micropipette aspiration analysis of RGG condensates under transmitted light. **a**, Time lapse transmitted (upper) and fluorescence (lower) images of an RGG condensate (adhered to a second pipette) under sequential ejection (−300 Pa, 0~20 s) and suction pressures (50 Pa, 20~40 s). **b**, Aspiration pressure (upper) and normalized aspiration length (lower) quantified from the transmitted light images. Gray lines: linear fits of the normalized aspiration length for each pressure step. **c**, *P*_asp_ of each step plotted against *V* (slopes of the gray lines in **b**). The black line represents a linear fit (slope: 660 ± 30 Pa·s, intercept: −100 ± 10 Pa, R^2^ = 0.985). Scale bars, 10 μm.

**Movie S1: Micropipette aspiration of an RGG condensate free from adhesion to glass surfaces.**

**Movie S2: Micropipette aspiration of an RGG condensate strongly adhered to a glass pipette.**

**Movie S3: Micropipette aspiration of an RGG condensate imaged with transmitted light.**

## Supplementary Table

| Liquid condensate | Viscosity (Pa·s) and method | Surface tension (mN/m) and method | Note |
| --- | --- | --- | --- |
| PGL-1<sup>4</sup> | ~1<br>FRAP | ~10<sup>-3</sup><br>Fusion | Order of magnitude estimates |
| LAF-1<sup>9</sup> | 23.4<br>SPT | 0.19<br>Fusion | Average over RNA concentrations |
| Whi3<sup>10</sup> | 15<br>FRAP and SPT | 5 x 10<sup>-5</sup><br>Fusion | Average over RNA concentrations |
| NPM1<sup>11</sup> | 0.74<br>FRAP and SPT | 8 x 10<sup>-4</sup><br>Fusion and Sessile drop |  |
| NPM1(*in vivo*)<sup>11</sup> | 37<br>FRAP | 4 x 10<sup>-4</sup><br>Fusion and Sessile drop |  |
| FIB1<sup>11</sup> | 100<br>FRAP and SPT | 1.23 x 10<sup>-3</sup><br>Fusion and Sessile drop |  |
| PGL-3<sup>12,13</sup> | 1 to 10<sup>4</sup><br>Dual-OT | ~4.5 x 10<sup>-3</sup><br>Dual-OT | The viscosity increased significantly over time |
| FUS<sup>12</sup> | 0.7 to 50<br>Dual-OT | ~3.1 x 10<sup>-3*</sup><br>Dual-OT | The viscosity increased significantly over time |
| Poly K<sup>14</sup> | 0.204<br>FRAP and SPT | 0.017<br>Fusion |  |
| Poly R<sup>14</sup> | 14.4<br>FRAP and SPT | 0.1<br>Fusion |  |
| [RGRGG]<sub>5</sub>-dT40<sup>15</sup> | 3<br>SPT and FCS | 0.8<br>Fusion-OT | Average over salt concentrations |
| Dextran-PEG<sup>16,17</sup> | 0.023~0.17<br>Viscometer | 0.01~0.1<br>Pendant drop | At compositions similar to sample used in this study |
| Olive oil<sup>18,19</sup> | 0.0741<br>Viscometer | 23.6<br>Pendent drop | All measured in water |
| Silicon oil<sup>20</sup> | 0.02<br>Viscometer | 36<br>Bubble contour |  |
| Mineral oil<sup>21</sup> | 7.75 x 10<sup>-3</sup><br>Viscometer | 49<br>Tensiometer |  |
| C<sub>16</sub>H<sub>34</sub><sup>22</sup> | 2.77 x 10<sup>-3</sup><br>Viscometer | 55.2<br>Tensiometer |  |
| C<sub>16</sub>H<sub>34</sub> with surfactants<sup>23</sup> | N.A. | Lowest to ~ 0.5<br>Tensiometer |  |
| C<sub>10</sub>H<sub>22</sub><sup>22</sup> | 9 x 10<sup>-4</sup><br>Viscometer | 53.2<br>Tensiometer |  |
| C<sub>6</sub>H<sub>14</sub><sup>22</sup> | 3.13 x 10<sup>-4</sup><br>Viscometer | 51.4<br>Tensiometer |  |

\* Surface tension value of FUS condensates was kindly provided by Dr. Frank Jülicher via email
**FRAP:** fluorescence recovery after photobleaching
**SPT:** single particle tracking
**Fusion:** first, a ratio of surface tension to viscosity was estimated from the fusion kinetics between two condensates. Then, a separate measure of viscosity was used to calculate surface tension values. Fusion experiments carried out using optical traps were noted as Fusion-OT.
**Sessile drop:** a prism was used to image condensates of various sizes from the side. The shape is determined by surface tension and gravity of the condensate. A separate measure of condensate density was used to extract surface tension.
**Dual-OT:** Dual optical traps were used to periodically stretch a condensate via two bead-handles. Viscoelasticity of the condensate was measured via the phase delay of the strain relative to the stress. Surface tension was estimated from the elasticity of condensates.

